# The gut microbiome mediates adaptation to scarce food in Coleoptera

**DOI:** 10.1101/2023.05.12.540564

**Authors:** Oana Teodora Moldovan, Alyssa A. Carrell, Paul Adrian Bulzu, Erika Levei, Ruxandra Bucur, Cristian Sitar, Luchiana Faur, Ionu□ Cornel Mirea, Marin □enilă, Oana Cadar, Mircea Podar

**Affiliations:** Emil Racovita Institute of Speleology, Cluj-Napoca Department, Clinicilor 5, 400006 Cluj-Napoca, Romania; Romanian Institute of Science and Technology, V. Fulicea 3, 400022 Cluj-Napoca, Romania; Centro Nacional de Investigación sobre la Evolución Humana, CENIEH, Paseo Sierra de Atapuerca 3, 09002 Burgos, Spain; Biosciences Division, Oak Ridge National Laboratory, Oak Ridge TN 37831, USA; Department of Aquatic Microbial Ecology, Institute of Hydrobiology, Biology Centre of the Academy of Sciences of the Czech Republic, 370 05 České Budějovice, Czech Republic; Research Institute for Analytical Instrumentation subsidiary, National Institute of Research and Development for Optoelectronics INOE 2000, Donath 67, 400293 Cluj-Napoca, Romania; Zoological Museum, Babe□ Bolyai University, Clinicilor 5, 400006 Cluj-Napoca, Romania; Emil Racovita Institute of Speleology, 13 Septembrie 13, 050711 Bucharest, Romania

**Keywords:** cave beetles, Carpathians, gut microbiome, sediments, adaptation

## Abstract

Beetles are ubiquitous cave invertebrates worldwide that adapted to scarce subterranean resources when they colonized caves. Here, we investigated the potential role of gut microbiota in the adaptation of beetles to caves from different climatic regions of the Carpathians. The beetles’ microbiota was host-specific, reflecting phylogenetic and nutritional adaptation. The microbial community structure further resolved conspecific beetles by caves suggesting microbiota-host coevolution and influences by local environmental factors. The detritivore species hosted a variety of bacteria known to decompose and ferment organic matter, suggesting turnover and host cooperative digestion of the sedimentary microbiota and allochthonous-derived nutrients. The cave Carabidae, with strong mandibulae adapted to predation and scavenging of animal and plant remains, had distinct microbiota dominated by symbiotic lineages *Spiroplasma* or *Wolbachia*. All beetles had relatively high levels of fermentative *Carnobacterium* and *Vagococcus* involved in lipid accumulation and a reduction of metabolic activity, and both features characterize adaptation to caves.

## Main

For centuries, naturalists have been intrigued by the long-legged eyeless beetles, transparent fishes and crustaceans that inhabit caves, leading to the science of biospeleology^1^. Caves are unique environments with no light, relatively constant climate, short food chains, and serve as a hub for research on adaptation, survival under extreme conditions, paleoclimate/ paleoenvironments, and even biomedicine^2,3,4^. Subterranean fauna is primarily represented by invertebrates, with Coleoptera (beetles) as the dominant non-aquatic group in caves, calcretes, or lava tubes^5^. Morphologically, beetles present various degrees of cave adaptation, culminating with slender bodies and extremely long legs and antennae. Regardless of the shape of the body and appendices, the cave-adapted (troglobiont) beetles are all eyeless or with reduced and depigmented eyes. Presumably originating from the so-called cryptic habitats (soil, under rocks or mosses^6^), the colonization of caves by beetles can be traced back to Oligocene^7^ and all species are endemic, often time to individual caves.

Caves have long been assumed to be nutrient depleted with their fauna dependent on surface food input. However recently, the discovery of a large diversity of microbes, that thrive in every subterranean biotope, many of them uncultured, suggests that the productivity and nutrient cycles in caves are more complex (e.g.^8.9^) and may sustain the local fauna. The gut microbiome of insects impacts their development, ecology, and evolution^10^ and contribute to the digestion and biosynthesis of essential metabolites^11,12^. Cave beetles are usually bigger than their surface relatives, lay one large egg, and have reduced developmental stages with a single non-feeding larval stage^13^. Such characteristics reflect cave adaptation and are likely linked to the gut microbiome composition and function^6^. Unlike the numerous studies on other insect microbiomes, only two studies on cave beetles have been published so far, revealing diverse bacterial communities in the guts of *Cansiliella* and *Neobathyscia*, both members of the Leptodirini tribe (family Leiodidae)^14,15^. Paoletti et al.^14^ sequenced cultivated bacteria of the midgut and found similar, but genetically diverged, bacteria common to animals’ digestive systems. Latella et al.^15^ characterized the gut of *Neobathyscia* using the PCR-DGGE technique and identified potential nitrogen-fixing bacteria and microbes with potential roles in the detoxification of poisonous wood compounds (such as tannins). Together these papers suggest the involvement of the gut microbiome in the evolution and adaptation of beetles to caves but the food sources and adaptation to low nutrient resources in caves remain unclear. Further, the interspecific regional variability of the gut microbiome remains to be elucidated. To better understand the evolution and adaptation of beetles to caves, we must characterize the role of cave location and food sources in shaping the gut microbiome of cave beetles.

Here, we utilized next-generation sequencing (NGS) to examine the diversity of the gut microbiota in seven endemic species of beetles, across seasons and from caves in different climate areas (**Figs. 1 and S1**). The species belong to two different tribes with supposed different food regimes, based on their mandibles’ morphology, the detritivores (Leptodirini, Leiodidae) and carnivores (Trechini, Carabidae). The Leptodirini species we selected are at different levels of cave adaptations. We hypothesized that in an environment with scarce food resources, the gut microbiota would co-evolve with its host and potentially serve as a proxy for the nutritional regimes in beetles with different degrees of cave adaptation. It will also differ among species and regions. The gut microbiome will also reveal if cave beetles depend exclusively on allochthonous, originating from the surface and seasonally variable, nutritional sources or use also autochthonous sources.

**Fig. 1:**
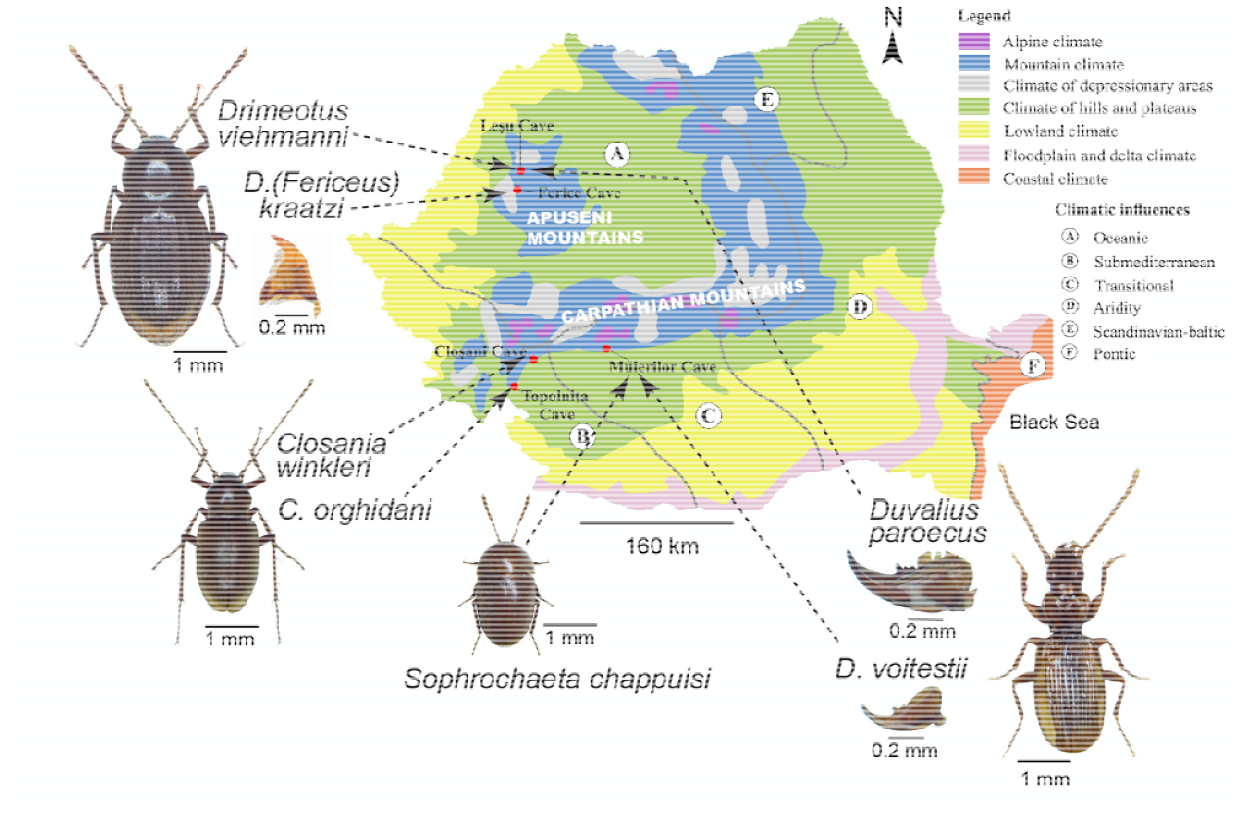
The sampled cave species and sites in the Carpathian Mountains. Caves from where representatives of Coleoptera were analyzed for the gut microbiome are shown within the climatic zonation and influences (modified after ^49^). The mandibles of a representative detritivore ((*Drimeotus (Fericeus) kraatzi*) and of the two *Duvalius* species are also shown.

## Results

### Seasonality, mineralogy, and geochemistry of cave environments

Expectedly, the general climate parameters were quite stable across seasons and caves, with less than 3 degrees of temperature variation at any station between summer and winter (8-13° C actual values across all caves; **Table S2**). Similarly, the air relative humidity values ranged between 70s-90s%, with lower values in the late spring-summer. Carbon dioxide levels were consistently higher in late summer in all caves due to water dripping in bringing more dissolved CO_2_ from the soils above. The substrates at the sampling stations varied significantly across stations and caves, with diverse mineralogical associations of silicates (quartz, clay minerals, albite), carbonates (calcite), and phosphates (hydroxyapatite and fluorapatite), in different proportions (**Table S3, Fig. S2 B**). Not surprisingly, there were also geochemical variations across the caves, stations, and sampling time (**Table S4**). Because our goal was to collect the sediments at the actual location where the beetles were found, some of the variations reflect the spatial heterogeneity at those stations (**Fig. S2 A**). Among the most important elements we considered here were nitrogen and carbon, both related to cave microbial productivity and nutrient sources for detritivores. Total nitrogen content was very low (under the detection limit) in most of the samples from Ferice, Closani, and Topolnita but reached over 0.5% in Lesu and Muieri, with the highest values in the spring-summer. The total carbon (TC) on the other hand varied considerably between and within caves, from <0.01% in Closani, <1% in Ferice, 2% in Topolnita, 5% in Lesu, to >10% in Muieri. Phosphate content was the highest in Muieri (0.5%), while total sulfur ranged from 35 to >700 mg/kg. Ca, Al, and Fe were major components of each sample, reflecting the mineralogic composition. The concentration of other elements was generally low and varied between stations and caves.

### The sediment microbiota

Across the 23 sediment samples collected from the five caves over multiple seasons, we identified 968 genus level taxa, representing 8 phyla of Archaea and 45 of Bacteria. On average, the relative abundance in each cave was the highest for Proteobacteria (35%), Acidobacterota (12%) and Actinobacterota (10%), in line with their general distribution in other soils and sediments. However, some taxa with few or no cultured representatives reached appreciable levels at some stations. Among those, GAL15, a bacterial candidate phylum previously identified in other subsurface environments but for which no cultures or even inferred physiological inferences have been reported, reached considerable levels (up to 26% in Topolnita, 35% in Closani and even 65% in Muieri). Bacteria that are primarily symbionts or parasites included *Patescibacteria* (up to 21% in Lesu, 30-35% in Ferice and Topolnita and 55% in Muieri), and *Dependentiae* (12% in Muieri). Among the Patescibacteria, the most represented were *Saccharimonadia* (TM7), *Parcubacteria, Microgenomatia* and *Berkelbacteria* (**Fig. 2**). *Nitrospirota*, a chemolithoautotrophic nitrite-oxidizing group, was also relatively abundant (12% in Lesu) and may contribute to fixing molecular nitrogen in the cave environment. Overall composition of the sediment microbiota across all sampled sites was quite similar, with 93% of all genus level taxa with at least 0.05% average abundance being present in all caves. Alpha diversity analysis revealed however dissimilarity across caves in community richness (Faith’s phylogenetic distance, Kruskal-Wallis difference of means H=27, p=1e^-5^), evenness (Pielou’s, H=15, p=0.003) and diversity (Shannon’s, H=18, p=0.001), differences supported also by pairwise Kruskal-Wallis tests for few but not all caves (e.g. Faith’s Ferice-Lesu or Topolnita-Muieri H=10.7 q=0.003). The communities also partially separated by cave following a principal coordinate analysis (PCoA) based on Bray Curtis dissimilarity (**Fig. 2 B**). Seasonality of sampling did not contribute to the variation, but we observed some stations effects in Lesu, Ferice, and Muieri that could not be linked to other measured parameters.

**Fig. 2:**
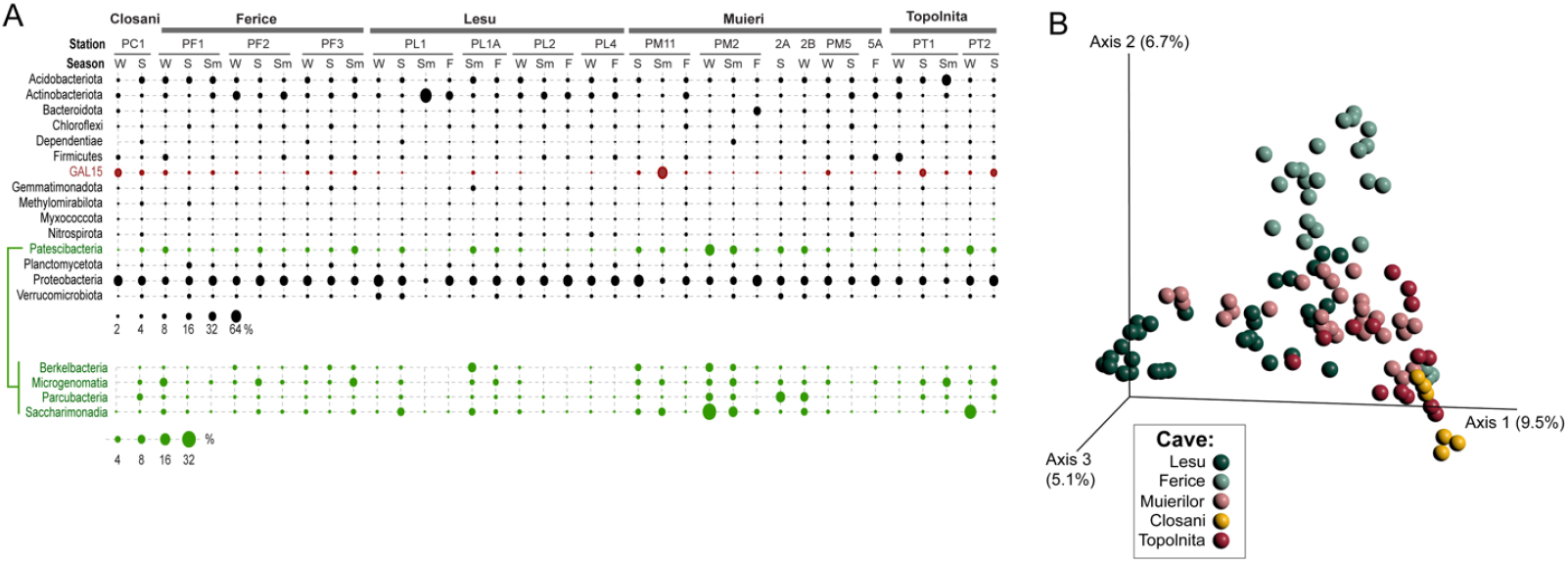
The distribution and abundances of bacteria phyla in the studied cave sediments. **A.** Average relative abundance of top represented phyla in cave sediments across stations and seasons (winter, spring, summer, fall). B. Principal component projection Bray Curtis dissimilarity distances between cave sediment microbiotas.

### Cave beetles gut microbiota

The ASVs obtained from the gut of the 102 collected cave Coleoptera individuals were mapped to 969 genus-level taxa, of which 160 were present at a relative abundance of over 0.1% in at least one individual. Unlike sediment samples matching their locations, specific community characteristics were associated with the different species of beetles (**Fig. 3 A**). Among the Leptodirini, the microbiota of *Sophrochaeta chappuisi*, the least cave-adapted species, was distinct in terms of alpha diversity from that of any other sampled coleopteran host (Shannon’s diversity pairwise K-W corrected tests ranged from q=3e^-4^-3e^-6^). The closely related, more troglomorphic *Drimeotus* and *D*. (*Fericeus)*, while originating from caves with distinct sedimentary microbiota (Lesu and Ferice, respectively) were indistinguishable (Shannon’s K-W q=0.91). A similar result was obtained for the most cave adapted detritivores Leptodirini, *Closania winkleri* and *C. orghidani* from Topolnita and Closani, respectively. The microbiota diversity of the suspected predatory *Duvalius* was distinct from that of any detritivore, including those they cohabitate with (Shannon’s K-W tests q=2-4e^-4^). Beta diversity analysis and PERMANOVA pairwise testing using Bray-Curtis distances also supported clustering of the coleopteran microbiota based on both troglomorphy and foraging/diet and are distinct from sedimentary communities (**Fig. 3 B**).

**Fig. 3:**
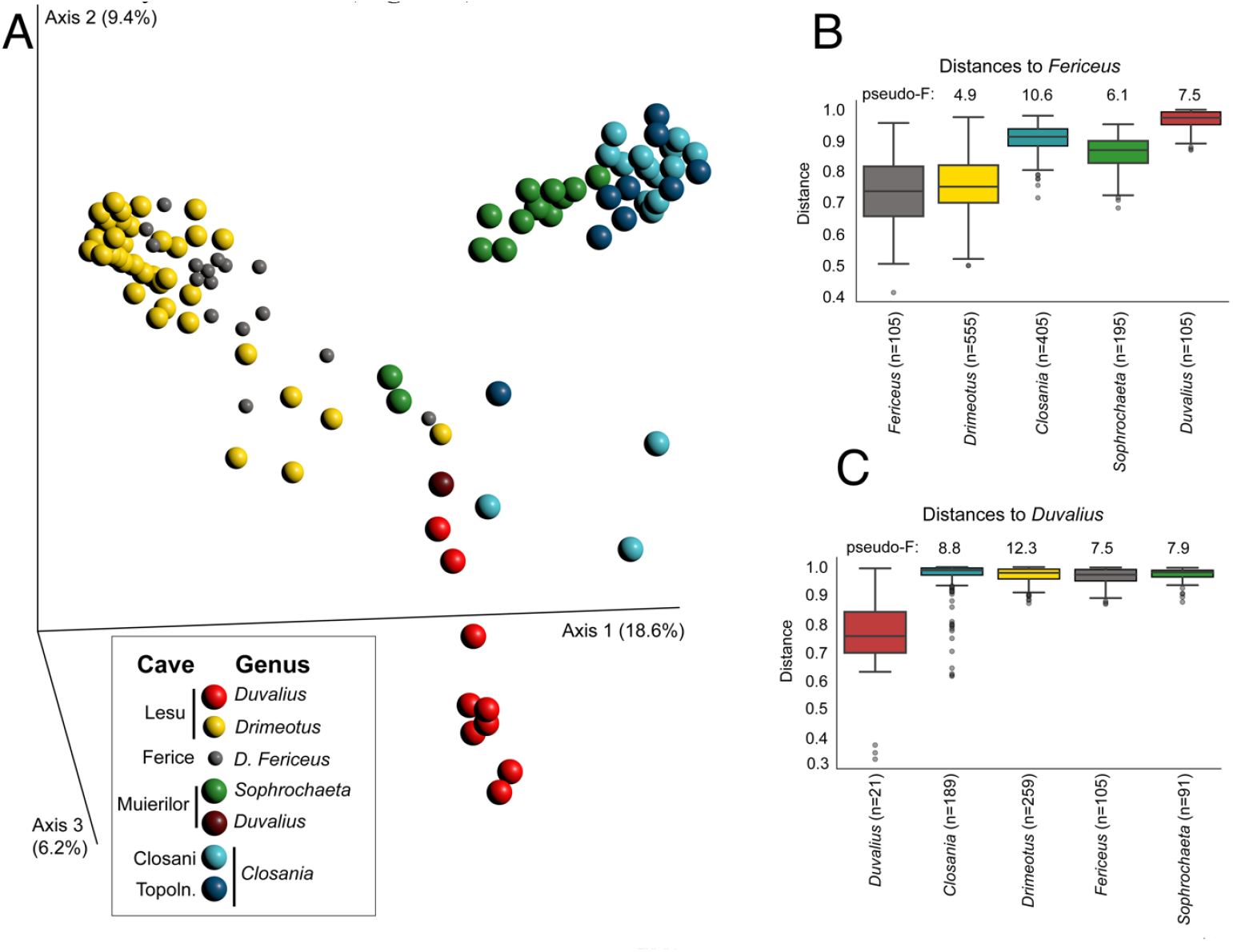
The gut microbiome of the studied cave Coleoptera. **A**. Principal component projection Bray Curtis dissimilarity distances between the gut microbiome of the different species; **B**. and **C**. Pairwise PERMANOVA between gut communities with 999 permutations, q = 0.001 based on BC distances, relative to *Drimeotus (Fericeus)* (B) and *Duvalius* from Muieri (C).

Overall, the gut microbiota of all detritivorous *Leptodirini* was dominated by the Firmicutes (on average, 41-55% across the various species and caves), followed by Bacteroidota (21% on average), Proteobacteria and Actinobacteriota (on average 13% and 12%, respectively; Fig. S3). Such general taxonomic distribution resembles the microbiota of many other animals, from mammals to insects. While there were individual beetle and species-level variations in relative abundance, overall, the general microbiota was remarkably conserved across this coleopteran group, even at the microbial genus level (**Fig. 4 A**). Among the most relatively abundant were Cand. *Soleaferrea, Tyzzerella* and *Vagococcus* from the Firmicutes, *Dysgonomonas* (Bacteroidota), *Arthrobacter* (Actionobacteriota) and *Ignatzschineria* (Gammaproteobacteria). The gut community of the *Duvalius paroecus* population on the other hand, is strikingly different from those of the *Leptodirini*, being dominated by *Spiroplasma* and *Carnobacterium* from the Firmicutes along with two Gammaproteobacteria, *Rickettsiella* and an uncultured member of the family *Wohlfahrtiimonadaceae*. In the only individual of *Duvalius voitestii* that we were able to collect, the gut microbiota differs greatly from that of its congeneric relative or any other collected coleopteran. The dominant bacteria in *D. voitestii* were *Spiroplasma* (Firmicutes), *Enhydrobacter* (Gammaproteobacteria) and *Wolbachia* (Alphaproteobacteria). The presence of *Wolbachia* in this individual is unique among the collected beetles and it is possible that such a typically endosymbiotic bacterium colonizes the gut tissue and did the luminal contents. Other bacterial representatives that were identified at low relative abundance include genera observed in some of the other collected beetles or sediment samples including *Micrococcus, Acinetobacter, Staphylococcus* and *Pseudomonas*. However, because it was a unique specimen, it was not included in statistical analyses.

**Fig. 4:**
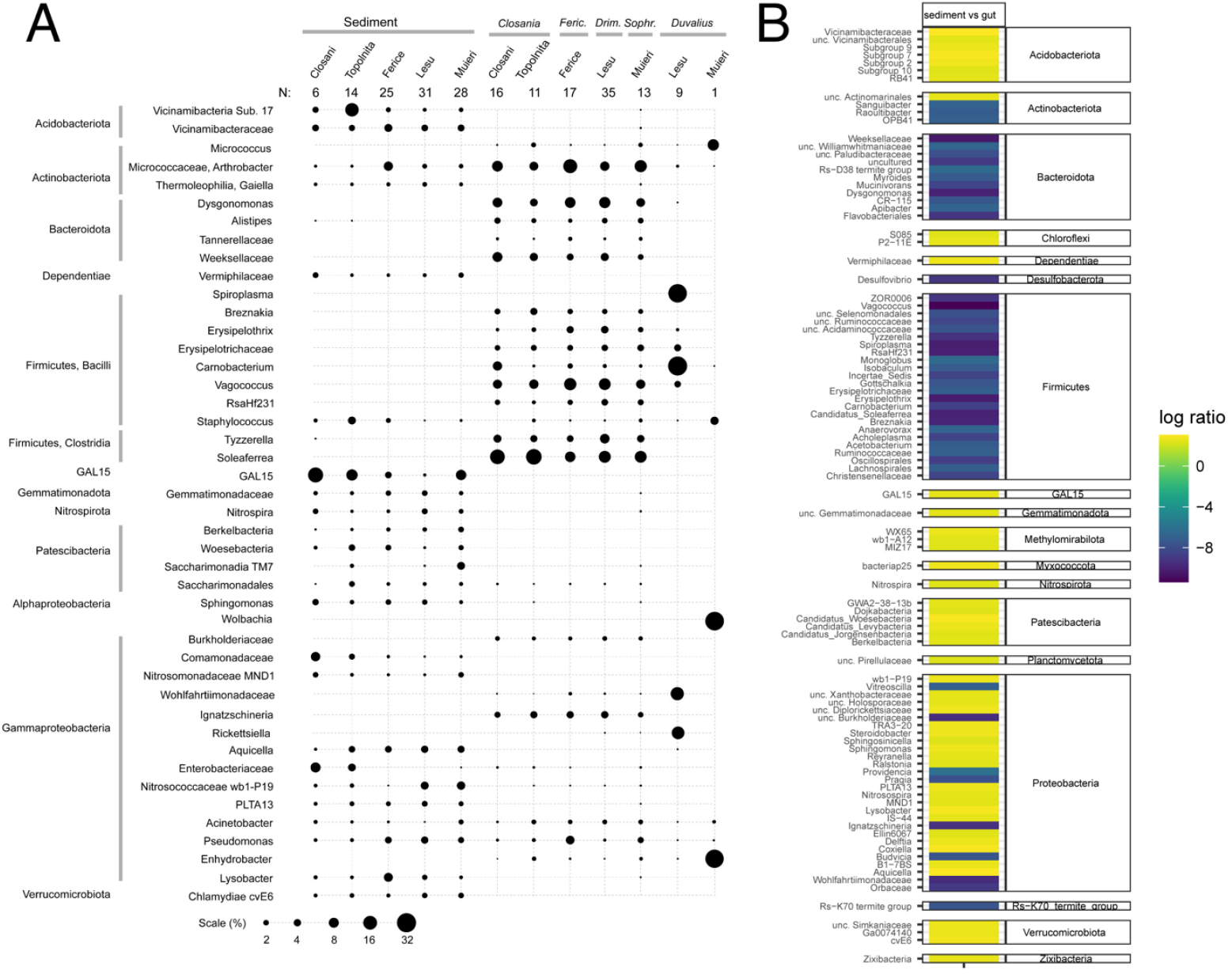
A comparative analysis between the sediment and gut microbiomes in the analyzed caves/species. **A**. Most relatively abundant genera in sediments and Coleoptera guts (averages). **B**. Differential ranking analysis of sediment versus beetle gut microbiota.

### Microbial specificity determinants across sediments and microbiomes

While the drastic differences between the cave sediments and the microbiota of the various species of Coleoptera readily revealed the relatively highly abundant microbes, we also aimed at detecting other microbes that were specific for various environments and species though at lower and variable relative abundance across individual samples. To achieve that, we applied differential ranking (DR) analysis following multinomial regression, an approach that enabled comparisons of compositional datasets that differ in absolute microbial load. When comparing the sediment environment with the coleopteran gut microbiome, regardless of cave or insect species, the sediment encompassed all the differentially enriched Acidobacteria and some typical environmental lineages (Chloroflexi, Dependetiae, GAL15, Nitrospirota, Patescibacteria, Planctomycetota, Verrucomicrobiota and others) (**Fig. 4 B**). On the other hand, all the differentially enriched lineages of Firmicutes and Bacteroidota were of coleopteran origin. They included the genera we observed by direct abundance analysis but revealed numerous others that, while at lower relative abundance, were strongly associated with the cave insect gut environment. Similar gut-characteristic lineages were identified among Actinobacterota and Proteobacteria. *Desulfovibrio* (Desulfobacterota), a sulfate reducer generally present in anoxic environments and in animal microbiota as well as an uncultured bacterium first identified in termites (Rs-K70) were also identified as coleopteran specific.

When comparing the various detritivore genera and species, few bacterial taxa were found to distinguish them (**Fig. 4 A**). Among them, *Arthrobacter* was generally absent from *Closania* individuals in both caves while *Vagococcus* was in a much higher abundance in the related *Drimeotus* and *D. (Fericeus)* than in the other genera. The distinctive *Duvalius* hosted, aside from the dominant *Spiroplasma* and *Carnobacterium*, harbored several other lineages that were absent in the detritivorous species including *Paenarthrobacter* (Actinobacteria), *Budvicia* and *Cedecea* (Enterobacteria).

## Discussion

Unlike sulfidic caves that are sustained by rich microbial chemolithoautotrophic production, microbial activity in oligotrophic carbonate and silicate caves relies primarily on allochthonous carbon (soluble and particulate, including plant biomass) from surface ecosystems, sourced by vadose and flowing water or introduced by macrofauna^16^. Several types of bacterial and archaea nitrifiers may support a nitrogen-based primary production in oligotrophic caves^8,17^ while diverse, metagenome-inferred carbon fixation pathways, suggest additional chemoautotrophic strategies may be at play^8,18^.

Among the core bacteria that may drive *in situ* primary production by nitrification, we identified abundant *Nitrospira* and *Nitrosomonadaceae* in all caves. We did not detect significant populations of archaeal chemolithoautotrophs. Instead, there were dominant chemoorganotrophs such as *Vicinamibacteria* (Acidobacterota) (especially in Topolnita) as well as cosmopolitan lineages of heterotrophic Actinobacterota, Firmicutes and Preoteobacteria, previously reported from other caves (e.g.^9,19^). Among them *Lysobacter* can decompose organic matter^20^ and provide fine particles for detritivores. Diverse lineages of putative symbiotic/parasitic bacteria (*Dependentiae, Berkelbacteria, Woesebacteria, Saccharibacteria*) were also abundant in all sediments. Interestingly, a candidate bacterial phylum, GAL15, was highly abundant in all caves and dominated the community at multiple stations in Closani and Muieri. Based on a handful of single-cell genomic and metagenomes, GAL15 bacteria have small genomes (∼1 Mbp) and are likely dependent on other microbes, with metabolic inferences pointing to adaptation to oligotrophic environments (i.e.^21^).

There was no direct correlation between the mineralogy, physical or chemical parameters of the cave sediments and their microbiota. Muieri is nevertheless distinct, as it has higher levels of carbon, nitrogen, phosphorus and sulfur and various trace elements, but with not distinct microbial diversity consequence. Similarly, seasonality had little effect on the on alpha and beta diversity across caves, although some specific taxa changed in relative abundance between individual sampling stations. In the caves we studied here, the substrate microbiota taxonomic structure is relatively uniform and stable within each cave and presents a baseline for comparisons an overall with the microbiomes of various beetle species whether detritivores or carnivores.

Across all caves and nutritional types, the beetle gut microbiota was drastically different than the sedimentary communities, which is inferred to represent an autochthonous nutritional source for the detritivore Leiodidae. Most of the abundant gut microbes were not present in the sediments, suggesting vertical transmission, from the surface. In animals, including insects, it has been shown that diet is directly correlated with gut microbial diversity, which increases from carnivores to omnivores and herbivores^22,23^. We observed the same in the cave beetles investigated here, the detritivore Leiodidae having a much more diverse gut microbiota than the predatory Carabidae.

Highly abundant bacteria in the Leiodidae gut microbiome are likely involved in lignocellulosic assimilation, including *Dysgonomonas, Clostridium, Tyzzerella*, and *Acinetobacter* (i.e.^24,25,26^), anaerobic fermentation (*Soleaferrea* and *Tyzzerella*^27^), and in decomposing carcasses (*Ignatzschineria, Carnobacterium, Rickettsia* and *Erysipelothrix)*. Some of these bacteria are carnivore biomarkers in Coleoptera and Orthoptera^28,29^ due to the presence of proteolytic and lipolytic enzymes. These dominant taxa define cave Leiodidae as opportunistic detritivores that can use different sources of organic matter available in caves, actively degrading and transforming them, including the sediments microbiota.

We found that the gut microbiome is host specific, with alpha and beta diversity reflecting beetle phylogenetic relationships, with the strong separation of Carabidae from Leiodidae and further of the phyletic lines in Leiodidae, down to species from the same geographic area. Even the species located in the same geographic area are relatively well separated, regardless of the degree of adaptation to cave life (e.g., *Sophrochaeta* vs *Closania*). Characteristic for the Leiodidae gut was the dominance of *Vagococcus, Dysgonomonas*, Candidatus *Soleaferrea, Carnobacterium* and uncultured Actinobacterota and Bacteroidota phylotypes. *Vagococcus* was previously reported in the *Neobathyscia* cave beetle^15^ and is co-dominant with *Lactococcus* during pre-diapause lipid accumulation in cabbage beetle (*Callophelus bowing*^30^).

The association in the gut microbiome of the studied cave beetles is unique with little overlap in the microbiomes associated with other groups of insects, not even with other Carabidae^31^. The dominance of *Vagococcus* in the detritivore Leptodirini and its presence in *Duvalius* (∼4%) is a marker for a metabolism inclined for lipid accumulation and a reduction of metabolic activity, like in diapausing surface insects. Diapause is an endocrine-mediated metabolic and developmental arrest^32^ and could play an essential role in the metabolism reduction of cave beetles. Colonizing caves, scarce in food sources, was not a drastic or sudden adaptation of the metabolism. It was merely a lengthening of a process similar but not identical to diapausing. Such a process was possibly mediated by a change in the dominant taxa in the gut microbiome, increasing the role of *Vagococcus*. Other adaptations in diapausing insects include physiological changes of the digestive system and fat accumulation^32,33^. The switch from diapause to reproduction and vice versa is mediated by the juvenile hormone (JH) that interacts with the circadian clock genes at the insect gut level^34^. Colonization of caves or smaller voids was made possible by the pre-adaptation of the gut microbiome, the prolongation of the torpor phase, which allows efficient use of scarce food, a slower metabolism, and longer life, known features of cave beetles^6^. Changes in the gut microbiome during colonization of the caves and adaptation to a different and low in nutrients food was not an abrupt process but one like in diapause, impacting the circadian clock genes. The lack of a circadian rhythm in the new subterranean environment was a hormone-mediated process that completed the adaptation to the underground lack of seasonality.

The gut microbiome in early spring (March) was somewhat more diverse (**Fig. S6**), potentially related to above soil microbiome seasonality^35^ and its effects underground. In the spring, when there is increased input of organic matter through the percolating water, the diversity of the gut microbiome at the individual level can increase. Spring is also a period of migration of cave beetles from the smaller voids into the cave, a more active period in the life of *Drimeotus viehmanni* in Lesu Cave^36^. Variations at the individual level in the *Duvalius* gut are speculative due to the low number of specimens we could analyze. In addition, some specimens had higher levels of obligate symbionts/parasites (*Spiroplasma, Ricketsiella* and *Wolbachia*), which impacts relative abundance levels of the other bacteria. However, it reveals, like in Leptodirini, the spring season separation from the summer and fall microbiomes. One specimen from Lesu in March and the May gut of Muieri were dominated by *Enhydrobacter* with cellulolytic potential^37^. The other most abundant classes in *Duvalius* gut, like Bacilli, Gammaproteobacteria, and Actinobacteria were previously documented in other insects^38^ but the originality we found was the presence of the dominant lactic acid *Carnobacterium* in the gut of *D. paroecus*. Lactic acid bacteria are known for increasing nutrient availability, producing antimicrobial substances, and improving disease resistance and antioxidative stress tolerance^39^. The other abundant genera define the gut of necrophagous beetles (*Wohlfahrtiimonas* and *Morganella*; ^40,41^), wood borer, *Ips* (*Tyzzerella*; ^42^), plants or phloem sap feeders (*Acinetobacter*; ^10^), or plant polymers feeders (*Lactococcus*; ^43^). Thus, *D. paroecus* gut microbiome points to an omnivore food regime. Their main food sources can be protein-and lipid-rich ones (possibly small cave invertebrates), carcasses, and even wood-related sources from the surface. On the contrary, *D. voitestii* has a completely different place in the cave food web with the gut microbiome associated only with plant or wood consumption. *Duvalius voitestii* fine and small-tooths mandible shows the adaptation to a lignivore food regime, and the analyzed individual is not an exception as the mandible shape is a genetic-conserved feature. A more productive region in a different climatic Carpathian region may have driven a niche partitioning between ancestors of *D. voitestii*, with the actual cave species adapted to the use of rich organic input from the surface. At present, *Duvalius* was empirically considered only in the frame of a predation-predator paradigm.

However, *Duvalius* can introduce microbes (i.e., *Lactococcus*) in the carrion or wood fragments, thus accelerating nutrient turnover and providing food for the detritivore invertebrates, as in necrophagous insects^44^. It offers, then, a completely different perspective on how food links might function in caves, in multiple directions (**Fig. 5**).

**Fig. 5:**
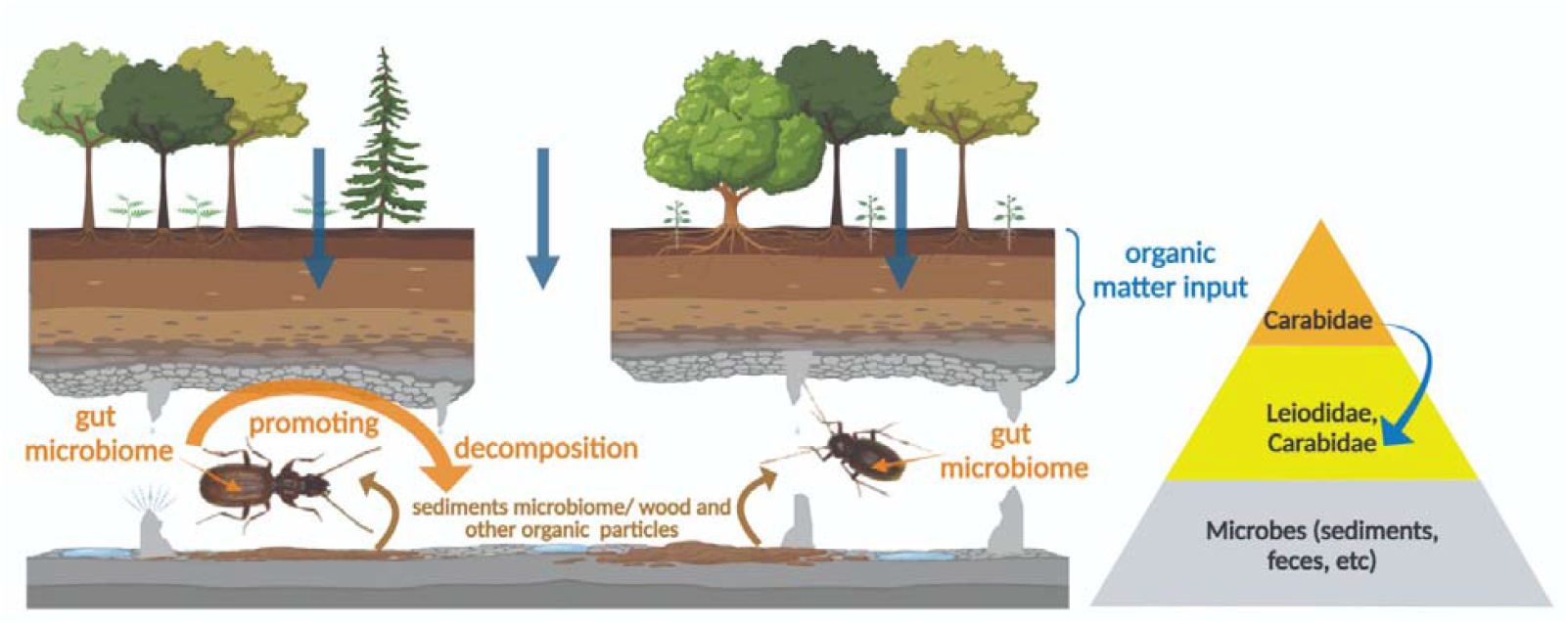
The simplified trophic relationships of the studied cave beetles and the cave microbiomes. The cave scheme with the food sources and microbiomes in caves and the role of the two-way omnivore carabid microbiome in the trophic chain, using the autochthonous and allochthonous resources and grinding the organic matter for the detritivore species.

The simplicity of the food webs in caves makes the cave inhabitants model organisms for the gut microbiome study in the context of adaptation to scarce food sources and limited competition/predation in other environments or as proxies for possible extraterrestrial habitats. Further, metagenomic analyses of cave beetles at different morphological stages of adaptation to life in caves will help to test these hypotheses and advance the understanding of the genetic mechanism of adaptation to a new environment.

## Material and Methods

### Locations and Sample collection

Caves beetles form small, endemic populations in some Romanian Carpathian caves. We selected five caves known to host diverse beetle species in specific areas (stations) in each cave (**Figs. 1 and S1**). The caves are hosted in karstic rocks (limestones, dolomites) and are in different geographic and climatic sub-regions of the Romanian Carpathians: Lesu and Ferice caves in the north-western Apuseni Mountains (oceanic climate), Clo ani and Topolni a (sub-Mediterranean) and Muierilor (transitional) in the south-western Carpathians (**Fig. 1**). Sampling was conducted seasonally in 2019: Late winter (March), spring (May), summer (August), and fall (November) (**Table S1**) and included sediment and climatic measurements (temperature, air humidity and air CO_2_ concentration). Further description of the caves and the sampling stations are provided in the **Table S1**. We selected cave-adapted, blind beetles with predicted different diets (detritivores and carnivores), at different stations, based on previous explorations. They belong to two different groups, *Leptodirini* (*Leiodidae*) and *Trechinae* (*Carabidae*). The detritivorous Leptodirini genera and species in this study have different degrees of cave adaptation (troglomorphic appearance, *sensu* ^45^). They range from the least troglomorphic *Sophrochaeta* with a small, round body and shortest antennae and legs, to the more elongated *Drimeotus* s. str. and *Drimeotus (Fericeus*), and the most troglomorphic *Closania*, with the longest antennae and legs (**Fig. 1**). *Drimeotus* is separated from *Closania* and *Sophrochaeta* in different phyletic lineages^46^. From the assumed carnivorous *Trechinae* (based on the mandibles morphology), we selected two species belonging to *Duvalius*, the only genus with cave representatives of this subfamily in the Carpathians. The two species show different degrees of troglomorphy (**Fig. 1**). *D. paroecus* has larger but unpigmented eyes and strong mandibles, while *D. voitestii* has completely reduced eyes and finer mandibles. We collected nine individuals of *D. paroecus*, while for *D. voitestii* only one specimen provided enough genetic material for sequencing. The collections were conducted in deep areas of the caves, with constant climate, and without attractors (baits). The individuals were collected with a fine brush or a sterile tweezer. They were stored in absolute ethanol, transported, and kept on ice until DNA extraction. Sediments for chemical, mineralogical and microbiological analyses were also sampled in triplicate from the same location where the beetles were found (Supplementary Material) and stored at -20°C until processing.

### Mineralogy and Geochemistry

Powdered X-ray diffraction analyses were performed on sediments to establish their mineralogy. Samples were analyzed with a Rigaku Ultima IV diffractometer in parallel beam geometry with CuKα radiation (wavelength 1.5406 Å). The XRD patterns were collected in 2Θ range between 5 to 80 with a speed of 2º/min and a step size of 0.02º. PDXL software from Rigaku, connected to the ICDD database, was used for phase identification. The quantitative determination was made using the RIR (Reference Intensity Ratio) methodology. The pH and electrical conductivity (EC) were measured in 1/5 (m/v) solid to water suspension with a Seven Excellence multimeter (Mettler Toledo, Greifensee, Switzerland). N and C were measured using a Flash 2000 CHNS/O analyzer (Thermo Fisher Scientific, Waltham, MA, USA). Major elements (Na, Mg, K, Ca, Al, Fe, P, and S) were measured using a 5300 Optima DV (Perkin Elmer, Waltham, Massachusetts, USA) inductively coupled plasma optical emission spectrometer. Trace elements (V, Cr, Mn, Co, Ni, Cu, Zn, As, Sr, Ba, La, Ce, and Pb) were measured with an Elan DRC II (Perkin Elmer, Waltham, Massachusetts, USA) inductively coupled plasma mass spectrometer, after *aqua regia* digestion.

### Gut isolation, DNA extraction and amplicon sequencing

The collected specimens were analyzed, identified, and dissected using standard entomological procedures under an Olympus SZX16 stereomicroscope, in sterile glass containers and using sterile utensils. The extracted guts were used immediately for total DNA extraction with the Quick-DNA Fecal/Soil Miniprep kit (Zymo Research, Irvine, CA), applying a 40-minute cell disruption with an Analog Genie Disruptor (Scientific Industries). DNA from sediment samples were extracted with the same kit and procedures. DNA quantification was performed with SpectraMax QuickDrop (Molecular Devices). SSU rRNA gene amplicon sequencing was conducted by a commercial company (Macrogen Europe, Amsterdam, Netherlands). Briefly, the V3-V4 hypervariable region of the bacterial and archaeal SSU rRNA gene was amplified using the universal primers 341F (5’CCTACGGGNGGCWGCAG3’) and 805R (5’GACTACHVGGGTATCTAATCC3’), according to Illumina’s 16S amplicon-based metagenomics sequencing protocol, followed by sequencing (2×300nt reads) on a MiSeq instrument (Illumina Inc, San Diego CA). Sequencing data generated in this study are available at the European Nucleotide Archive (ENA) under the study accession number PRJEB61400.

### Amplicon sequence processing, taxonomic and statistical analyses

Paired amplicon reads, demultiplexed based on samples, were imported into QIIME2 v.2021.2^47^. Quality based denoising, dereplicating, chimera filtration and amplicon sequence variant (ASV) identification were performed with the DADA2 plugin, applying trimming to remove primers and low quality read regions (--p-trim-left-f 17, -r 21, --p-trunc-len-f 260, -r 230). The ASVs were classified to different taxonomic levels based on the SILVA-v138 database. Standard QIIME2 workflows were used for determining alpha (Shannon’s diversity index, Faith’s Phylogenetic Diversity and Pielou’s Evenness) and beta diversity (Bray-Curtis dissimilarity and UniFrac distances), using rarefaction at 10,000 sequences per sample. The samples were analyzed collectively or split into specific groups to test hypotheses of the effects of environment (sediment vs gut microbiota, seasonality, cave, beetle species or feeding type) on diversity or composition using one-way ANOVA (Kruskal-Wallis H test) and pairwise PERMANOVA tests implemented in QIIME2. To visualize and explore the relationship between the samples based on various metadata, beta diversity distances were subjected to principal coordinates analyses (PCoA) and projected on Emperor plots that used the first three dimensions using QIIME2 View. Differential abundance analysis to identify microbes specific for environment (sediment versus beetle gut) or between types or species of beetles was performed using Songbird^48^ at the genus taxonomic level through QIIME 2 v2020.6. Briefly, data was filtered to include genera present in a minimum of 10 samples for environment comparison and genera present in a minimum of 5 samplers for the beetle species comparison. Relative differentials were estimated using multinomial regression with taxonomy collapsed to the genus level. Models were checked for fit with visualization of cross-validation and loss as well as validated against null models and used for differential ranking of taxa across compared datasets (e.g., sediment serving as reference in environment comparisons or *Duvalius* in beetle comparisons). Log-ratios of the differential were filtered for the top positive and negative log-ratios (top 50 positive and 50 negative for sediment vs gut and top 25 positive and negative for each beetle in the beetle species comparison) and displayed as heatmaps to track positive versus negative differences between the compared data types at genus level.

## Acknowledgments

We are grateful to our colleagues Marius Kenesz, Alexandru Petculescu and Răzvan Arghir that helped with the sampling and to Victor Fruth and Irina Atkinson from the Faculty of Geology and Geophysics, University of Bucharest, for the help in analyzing some of the sediment samples. The collection of samples was done under permit no. 7/2019 issued by the National Commission of the Speleological Heritage (Romania). This research was supported by a grant from the Ministry of Research, Innovation and Digitization, CNCS/CCCDI – UEFISCDI, project number 2/2019 (DARKFOOD) within PNCDI III, and the project EEA 126/2018 (KARSTHIVES2), contract no. 3/2019. AAC and MP were supported by the U.S. Department of Energy, Office of Science, Biological and Environmental Research and by grant R01DE024463 from the National Institute of Dental and Craniofacial Research of the US National Institutes of Health. Oak Ridge National Laboratory is managed by UT-Battelle, LLC, for the U.S. Department of Energy under contract DE-AC05-00OR22725. P-AB was supported by research grant no. 20-23718Y (Grant Agency of the Czech Republic).

## Notes

### Competing Interest Statement

The authors have declared no competing interest.

