## Supplemental material for "The gut microbiome mediates adaptation to scarce food in Coleoptera"

**Supplementary Material**

Sites description.

Apă din Valea Leșului Cave (Lesu), located at an altitude of 700 meters, is horizontal and populated by large hibernation colonies of bats. Station 1 (L1) is located on a sandy beach about 500 m from the entrance. Station L1A is about 20 m upstream from L1 in a lateral passage. Stations L2 and L4 are about 10 m upstream from L1A on the left side of the main gallery, in a small side passage (Fig. 1). Cave temperature varies between 8 and 11 °C, depending on the season, while the relative air humidity was > 80% in August and November and >90% in March and May (Table S1).

Ferice Cave (Ferice) is located at an altitude of 410 meters. The main passage is horizontal with a length of approximately 260 m, crossed by a low-flow stream. The cave is used by very few bats, especially in winter. Station F1 is about 60 m from the entrance. The other two stations (F2 and F3) are located ca. 100 and 200 m, relatively, from the entrance (Fig. 1). The air temperature was relatively constant throughout the year (between 11.4 and 12.7 °C), while the humidity was high (~ 95%) during November and the lowest in May (~80%) (Table S1).

Cloşani Cave (Closani), located at an absolute altitude of 433 meters, is a dry cave, 1100 m long, developed on two passages, and hosts occasional bat colonies. The two collecting and monitoring stations (C1 and C3) were located on the left side gallery, at about 150 and 250 m from the entrance (Fig. 1). The relatively constant temperature was also registered in this cave (11.2 °C in May and November and 12-13 °C in March and August). Air humidity was lower than 90% only in August (Table S1).

Topolnița Cave (Topolnita) consists of a network of passages of about 20 km. The sampling and monitoring stations are in the dry sector of the system, with access at an altitude of 429 meters. Significant bat colonies and guano deposits are in the cave near T1 station, located on sandy-clay substrate (Fig. 1). Station T2 is at a 50 m distance from T1 on a calcite floor. Station T3 is 50 m from station T2 and is characterized by a sandy substrate on the cave floor. In Topolnita, air temperature varied around 11.5 °C in August and November, with higher values (12-14 °C) in the two other months. Air humidity was >94%, except for May (~85%) (Table S1).

Muierilor de la Baia de Fier Cave (Muieri), located at an altitude of 650 meters, is developed on four levels. The sampling and monitoring stations were in the upper level (PM2) and the lower level (PM5 and PM11; Fig. 1). The temperature varied in this cave around 10 °C, reaching ~12 °C in August and March only in M2 (Table S1). In this cave, fossil bones and guano deposits led to the precipitation of phosphate minerals such as hydroxyapatite and fluorapatite and phosphate-related microorganisms (Haidău et al., 2022).

We avoided collecting the samples in places with bats or guano, in all caves.

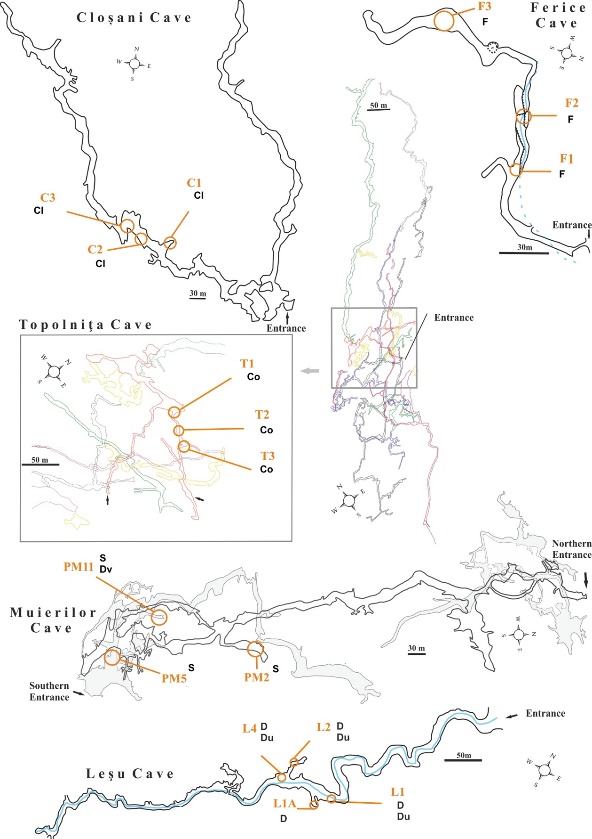

**Fig S1:** **The maps of the studied caves and the stations where sediment and beetles were sampled.** See Tables S1 and S2 for more information on the stations. Cl = *Closania winkleri*, Co = *Closania orghidani*, D = *Drimeotus viehmanni*, Du = *Duvalius procerus*, Dv = *Duvalius voitestii*, F = *Drimeotus (Fericeus) kraatzi*, S = *Sophrochaeta chappuisi*. Cloșani Cave map modified after Diaconu (1990); Ferice Cave map modified after Bleahu et al. (1976); Topolnița Cave map modified after Goran & Povară (2019); Muierilor Cave map modified after Mirea et al. (2021); Leșu Cave map modified after Rusu (1988).

A

|  | Dissimilarity |
| --- | --- |
| 12.0156 | 135.18 |
| 6.21875 | 135.18 |
| 6.21875 | 58.8755 |
| 2.5 | 58.8755 |
| 2.5 | 34.0032 |
| 1.5 | 34.0032 |
| 1.5 | 11.7855 |
| 1 | 11.7855 |
| 1 | 0 |
| 1 | 11.7855 |
| 2 | 11.7855 |
| 2 | 0 |
| 2 | 11.7855 |
| 1.5 | 11.7855 |
| 1.5 | 34.0032 |
| 3.5 | 34.0032 |
| 3.5 | 25.0159 |
| 3 | 25.0159 |
| 3 | 0 |
| 3 | 25.0159 |
| 4 | 25.0159 |
| 4 | 0 |
| 4 | 25.0159 |
| 3.5 | 25.0159 |
| 3.5 | 34.0032 |
| 2.5 | 34.0032 |
| 2.5 | 58.8755 |
| 9.9375 | 58.8755 |
| 9.9375 | 47.9132 |
| 7.25 | 47.9132 |
| 7.25 | 16.5757 |
| 5.75 | 16.5757 |
| 5.75 | 6.54125 |
| 5 | 6.54125 |
| 5 | 0 |
| 5 | 6.54125 |
| 6.5 | 6.54125 |
| 6.5 | 3.62943 |
| 6 | 3.62943 |
| 6 | 0 |
| 6 | 3.62943 |
| 7 | 3.62943 |
| 7 | 0 |
| 7 | 3.62943 |
| 6.5 | 3.62943 |
| 6.5 | 6.54125 |
| 5.75 | 6.54125 |
| 5.75 | 16.5757 |
| 8.75 | 16.5757 |
| 8.75 | 6.68248 |
| 8 | 6.68248 |
| 8 | 0 |
| 8 | 6.68248 |
| 9.5 | 6.68248 |
| 9.5 | 5.02365 |
| 9 | 5.02365 |
| 9 | 0 |
| 9 | 5.02365 |
| 10 | 5.02365 |
| 10 | 0 |
| 10 | 5.02365 |
| 9.5 | 5.02365 |
| 9.5 | 6.68248 |
| 8.75 | 6.68248 |
| 8.75 | 16.5757 |
| 7.25 | 16.5757 |
| 7.25 | 47.9132 |
| 12.625 | 47.9132 |
| 12.625 | 24.6713 |
| 11.5 | 24.6713 |
| 11.5 | 5.2274 |
| 11 | 5.2274 |
| 11 | 0 |
| 11 | 5.2274 |
| 12 | 5.2274 |
| 12 | 0 |
| 12 | 5.2274 |
| 11.5 | 5.2274 |
| 11.5 | 24.6713 |
| 13.75 | 24.6713 |
| 13.75 | 10.2805 |
| 13 | 10.2805 |
| 13 | 0 |
| 13 | 10.2805 |
| 14.5 | 10.2805 |
| 14.5 | 8.80241 |
| 14 | 8.80241 |
| 14 | 0 |
| 14 | 8.80241 |
| 15 | 8.80241 |
| 15 | 0 |
| 15 | 8.80241 |
| 14.5 | 8.80241 |
| 14.5 | 10.2805 |
| 13.75 | 10.2805 |
| 13.75 | 24.6713 |
| 12.625 | 24.6713 |
| 12.625 | 47.9132 |
| 9.9375 | 47.9132 |
| 9.9375 | 58.8755 |
| 6.21875 | 58.8755 |
| 6.21875 | 135.18 |
| 17.8125 | 135.18 |
| 17.8125 | 69.7685 |
| 16.5 | 69.7685 |
| 16.5 | 22.1091 |
| 16 | 22.1091 |
| 16 | 0 |
| 16 | 22.1091 |
| 17 | 22.1091 |
| 17 | 0 |
| 17 | 22.1091 |
| 16.5 | 22.1091 |
| 16.5 | 69.7685 |
| 19.125 | 69.7685 |
| 19.125 | 24.2827 |
| 18 | 24.2827 |
| 18 | 0 |
| 18 | 24.2827 |
| 20.25 | 24.2827 |
| 20.25 | 21.7565 |
| 19 | 21.7565 |
| 19 | 0 |
| 19 | 21.7565 |
| 21.5 | 21.7565 |
| 21.5 | 19.7197 |
| 20.5 | 19.7197 |
| 20.5 | 8.01275 |
| 20 | 8.01275 |
| 20 | 0 |
| 20 | 8.01275 |
| 21 | 8.01275 |
| 21 | 0 |
| 21 | 8.01275 |
| 20.5 | 8.01275 |
| 20.5 | 19.7197 |
| 22.5 | 19.7197 |
| 22.5 | 9.14309 |
| 22 | 9.14309 |
| 22 | 0 |
| 22 | 9.14309 |
| 23 | 9.14309 |
| 23 | 0 |
| 23 | 9.14309 |
| 22.5 | 9.14309 |
| 22.5 | 19.7197 |
| 21.5 | 19.7197 |
| 21.5 | 21.7565 |
| 20.25 | 21.7565 |
| 20.25 | 24.2827 |
| 19.125 | 24.2827 |
| 19.125 | 69.7685 |
| 17.8125 | 69.7685 |
| 17.8125 | 135.18 |
| 12.0156 | 135.18 |

|  | Dissimilarity |
| --- | --- |
| 6.21875 | 58.8755 |
| 2.5 | 58.8755 |
| 2.5 | 34.0032 |
| 1.5 | 34.0032 |
| 1.5 | 11.7855 |
| 1 | 11.7855 |
| 1 | 0 |
| 1 | 11.7855 |
| 2 | 11.7855 |
| 2 | 0 |
| 2 | 11.7855 |
| 1.5 | 11.7855 |
| 1.5 | 34.0032 |
| 3.5 | 34.0032 |
| 3.5 | 25.0159 |
| 3 | 25.0159 |
| 3 | 0 |
| 3 | 25.0159 |
| 4 | 25.0159 |
| 4 | 0 |
| 4 | 25.0159 |
| 3.5 | 25.0159 |
| 3.5 | 34.0032 |
| 2.5 | 34.0032 |
| 2.5 | 58.8755 |
| 9.9375 | 58.8755 |
| 9.9375 | 47.9132 |
| 7.25 | 47.9132 |
| 7.25 | 16.5757 |
| 5.75 | 16.5757 |
| 5.75 | 6.54125 |
| 5 | 6.54125 |
| 5 | 0 |
| 5 | 6.54125 |
| 6.5 | 6.54125 |
| 6.5 | 3.62943 |
| 6 | 3.62943 |
| 6 | 0 |
| 6 | 3.62943 |
| 7 | 3.62943 |
| 7 | 0 |
| 7 | 3.62943 |
| 6.5 | 3.62943 |
| 6.5 | 6.54125 |
| 5.75 | 6.54125 |
| 5.75 | 16.5757 |
| 8.75 | 16.5757 |
| 8.75 | 6.68248 |
| 8 | 6.68248 |
| 8 | 0 |
| 8 | 6.68248 |
| 9.5 | 6.68248 |
| 9.5 | 5.02365 |
| 9 | 5.02365 |
| 9 | 0 |
| 9 | 5.02365 |
| 10 | 5.02365 |
| 10 | 0 |
| 10 | 5.02365 |
| 9.5 | 5.02365 |
| 9.5 | 6.68248 |
| 8.75 | 6.68248 |
| 8.75 | 16.5757 |
| 7.25 | 16.5757 |
| 7.25 | 47.9132 |
| 12.625 | 47.9132 |
| 12.625 | 24.6713 |
| 11.5 | 24.6713 |
| 11.5 | 5.2274 |
| 11 | 5.2274 |
| 11 | 0 |
| 11 | 5.2274 |
| 12 | 5.2274 |
| 12 | 0 |
| 12 | 5.2274 |
| 11.5 | 5.2274 |
| 11.5 | 24.6713 |
| 13.75 | 24.6713 |
| 13.75 | 10.2805 |
| 13 | 10.2805 |
| 13 | 0 |
| 13 | 10.2805 |
| 14.5 | 10.2805 |
| 14.5 | 8.80241 |
| 14 | 8.80241 |
| 14 | 0 |
| 14 | 8.80241 |
| 15 | 8.80241 |
| 15 | 0 |
| 15 | 8.80241 |
| 14.5 | 8.80241 |
| 14.5 | 10.2805 |
| 13.75 | 10.2805 |
| 13.75 | 24.6713 |
| 12.625 | 24.6713 |
| 12.625 | 47.9132 |
| 9.9375 | 47.9132 |
| 9.9375 | 58.8755 |
| 6.21875 | 58.8755 |

|  | Dissimilarity |
| --- | --- |
| 17.8125 | 69.7685 |
| 16.5 | 69.7685 |
| 16.5 | 22.1091 |
| 16 | 22.1091 |
| 16 | 0 |
| 16 | 22.1091 |
| 17 | 22.1091 |
| 17 | 0 |
| 17 | 22.1091 |
| 16.5 | 22.1091 |
| 16.5 | 69.7685 |
| 19.125 | 69.7685 |
| 19.125 | 24.2827 |
| 18 | 24.2827 |
| 18 | 0 |
| 18 | 24.2827 |
| 20.25 | 24.2827 |
| 20.25 | 21.7565 |
| 19 | 21.7565 |
| 19 | 0 |
| 19 | 21.7565 |
| 21.5 | 21.7565 |
| 21.5 | 19.7197 |
| 20.5 | 19.7197 |
| 20.5 | 8.01275 |
| 20 | 8.01275 |
| 20 | 0 |
| 20 | 8.01275 |
| 21 | 8.01275 |
| 21 | 0 |
| 21 | 8.01275 |
| 20.5 | 8.01275 |
| 20.5 | 19.7197 |
| 22.5 | 19.7197 |
| 22.5 | 9.14309 |
| 22 | 9.14309 |
| 22 | 0 |
| 22 | 9.14309 |
| 23 | 9.14309 |
| 23 | 0 |
| 23 | 9.14309 |
| 22.5 | 9.14309 |
| 22.5 | 19.7197 |
| 21.5 | 19.7197 |
| 21.5 | 21.7565 |
| 20.25 | 21.7565 |
| 20.25 | 24.2827 |
| 19.125 | 24.2827 |
| 19.125 | 69.7685 |
| 17.8125 | 69.7685 |

|  | Dissimilarity |
| --- | --- |
| 0 | 102.474 |
| 24 | 102.474 |

|  | Dissimilarity |
| --- | --- |
| 1 | 0 |
| 2 | 0 |
| 3 | 0 |
| 4 | 0 |
| 5 | 0 |
| 6 | 0 |
| 7 | 0 |
| 8 | 0 |
| 9 | 0 |
| 10 | 0 |
| 11 | 0 |
| 12 | 0 |
| 13 | 0 |
| 14 | 0 |
| 15 | 0 |
| 16 | 0 |
| 17 | 0 |
| 18 | 0 |
| 19 | 0 |
| 20 | 0 |
| 21 | 0 |
| 22 | 0 |
| 23 | 0 |

B

|  | Dissimilarity |
| --- | --- |
| 5.57812 | 23.8085 |
| 1 | 23.8085 |
| 1 | 0 |
| 1 | 23.8085 |
| 10.1562 | 23.8085 |
| 10.1562 | 19.5958 |
| 6.8125 | 19.5958 |
| 6.8125 | 10.3936 |
| 4.6875 | 10.3936 |
| 4.6875 | 6.19309 |
| 2.875 | 6.19309 |
| 2.875 | 2.29727 |
| 2 | 2.29727 |
| 2 | 0 |
| 2 | 2.29727 |
| 3.75 | 2.29727 |
| 3.75 | 1.17189 |
| 3 | 1.17189 |
| 3 | 0 |
| 3 | 1.17189 |
| 4.5 | 1.17189 |
| 4.5 | 0.272232 |
| 4 | 0.272232 |
| 4 | 0 |
| 4 | 0.272232 |
| 5 | 0.272232 |
| 5 | 0 |
| 5 | 0.272232 |
| 4.5 | 0.272232 |
| 4.5 | 1.17189 |
| 3.75 | 1.17189 |
| 3.75 | 2.29727 |
| 2.875 | 2.29727 |
| 2.875 | 6.19309 |
| 6.5 | 6.19309 |
| 6.5 | 3.39071 |
| 6 | 3.39071 |
| 6 | 0 |
| 6 | 3.39071 |
| 7 | 3.39071 |
| 7 | 0 |
| 7 | 3.39071 |
| 6.5 | 3.39071 |
| 6.5 | 6.19309 |
| 4.6875 | 6.19309 |
| 4.6875 | 10.3936 |
| 8.9375 | 10.3936 |
| 8.9375 | 9.45632 |
| 8 | 9.45632 |
| 8 | 0 |
| 8 | 9.45632 |
| 9.875 | 9.45632 |
| 9.875 | 5.84597 |
| 9 | 5.84597 |
| 9 | 0 |
| 9 | 5.84597 |
| 10.75 | 5.84597 |
| 10.75 | 1.91475 |
| 10 | 1.91475 |
| 10 | 0 |
| 10 | 1.91475 |
| 11.5 | 1.91475 |
| 11.5 | 1.20859 |
| 11 | 1.20859 |
| 11 | 0 |
| 11 | 1.20859 |
| 12 | 1.20859 |
| 12 | 0 |
| 12 | 1.20859 |
| 11.5 | 1.20859 |
| 11.5 | 1.91475 |
| 10.75 | 1.91475 |
| 10.75 | 5.84597 |
| 9.875 | 5.84597 |
| 9.875 | 9.45632 |
| 8.9375 | 9.45632 |
| 8.9375 | 10.3936 |
| 6.8125 | 10.3936 |
| 6.8125 | 19.5958 |
| 13.5 | 19.5958 |
| 13.5 | 12.4513 |
| 13 | 12.4513 |
| 13 | 0 |
| 13 | 12.4513 |
| 14 | 12.4513 |
| 14 | 0 |
| 14 | 12.4513 |
| 13.5 | 12.4513 |
| 13.5 | 19.5958 |
| 10.1562 | 19.5958 |
| 10.1562 | 23.8085 |
| 5.57812 | 23.8085 |

|  | Dissimilarity |
| --- | --- |
| 1 | 0 |

|  | Dissimilarity |
| --- | --- |
| 4.6875 | 6.19309 |
| 2.875 | 6.19309 |
| 2.875 | 2.29727 |
| 2 | 2.29727 |
| 2 | 0 |
| 2 | 2.29727 |
| 3.75 | 2.29727 |
| 3.75 | 1.17189 |
| 3 | 1.17189 |
| 3 | 0 |
| 3 | 1.17189 |
| 4.5 | 1.17189 |
| 4.5 | 0.272232 |
| 4 | 0.272232 |
| 4 | 0 |
| 4 | 0.272232 |
| 5 | 0.272232 |
| 5 | 0 |
| 5 | 0.272232 |
| 4.5 | 0.272232 |
| 4.5 | 1.17189 |
| 3.75 | 1.17189 |
| 3.75 | 2.29727 |
| 2.875 | 2.29727 |
| 2.875 | 6.19309 |
| 6.5 | 6.19309 |
| 6.5 | 3.39071 |
| 6 | 3.39071 |
| 6 | 0 |
| 6 | 3.39071 |
| 7 | 3.39071 |
| 7 | 0 |
| 7 | 3.39071 |
| 6.5 | 3.39071 |
| 6.5 | 6.19309 |
| 4.6875 | 6.19309 |

|  | Dissimilarity |
| --- | --- |
| 8.9375 | 9.45632 |
| 8 | 9.45632 |
| 8 | 0 |
| 8 | 9.45632 |
| 9.875 | 9.45632 |
| 9.875 | 5.84597 |
| 9 | 5.84597 |
| 9 | 0 |
| 9 | 5.84597 |
| 10.75 | 5.84597 |
| 10.75 | 1.91475 |
| 10 | 1.91475 |
| 10 | 0 |
| 10 | 1.91475 |
| 11.5 | 1.91475 |
| 11.5 | 1.20859 |
| 11 | 1.20859 |
| 11 | 0 |
| 11 | 1.20859 |
| 12 | 1.20859 |
| 12 | 0 |
| 12 | 1.20859 |
| 11.5 | 1.20859 |
| 11.5 | 1.91475 |
| 10.75 | 1.91475 |
| 10.75 | 5.84597 |
| 9.875 | 5.84597 |
| 9.875 | 9.45632 |
| 8.9375 | 9.45632 |

|  | Dissimilarity |
| --- | --- |
| 13 | 0 |

|  | Dissimilarity |
| --- | --- |
| 14 | 0 |

|  | Dissimilarity |
| --- | --- |
| 0 | 9.92494 |
| 15 | 9.92494 |

|  | Dissimilarity |
| --- | --- |
| 1 | 0 |
| 2 | 0 |
| 3 | 0 |
| 4 | 0 |
| 5 | 0 |
| 6 | 0 |
| 7 | 0 |
| 8 | 0 |
| 9 | 0 |
| 10 | 0 |
| 11 | 0 |
| 12 | 0 |
| 13 | 0 |
| 14 | 0 |

**Fig. S2:** **The physicochemical and mineralogical dissimilarity between the sediments of the five studied caves.** **A.** Sediments from Muieri are physicochemically significantly separated from the other caves, with a single sample in station 1A Lesu as an exception. **B.** Mineralogy separated the northern (L and F) from the southern caves (C, T, and M) at low significance, with the exceptions of F3 (high content in chlorite) and T1 (high content in dolomite).

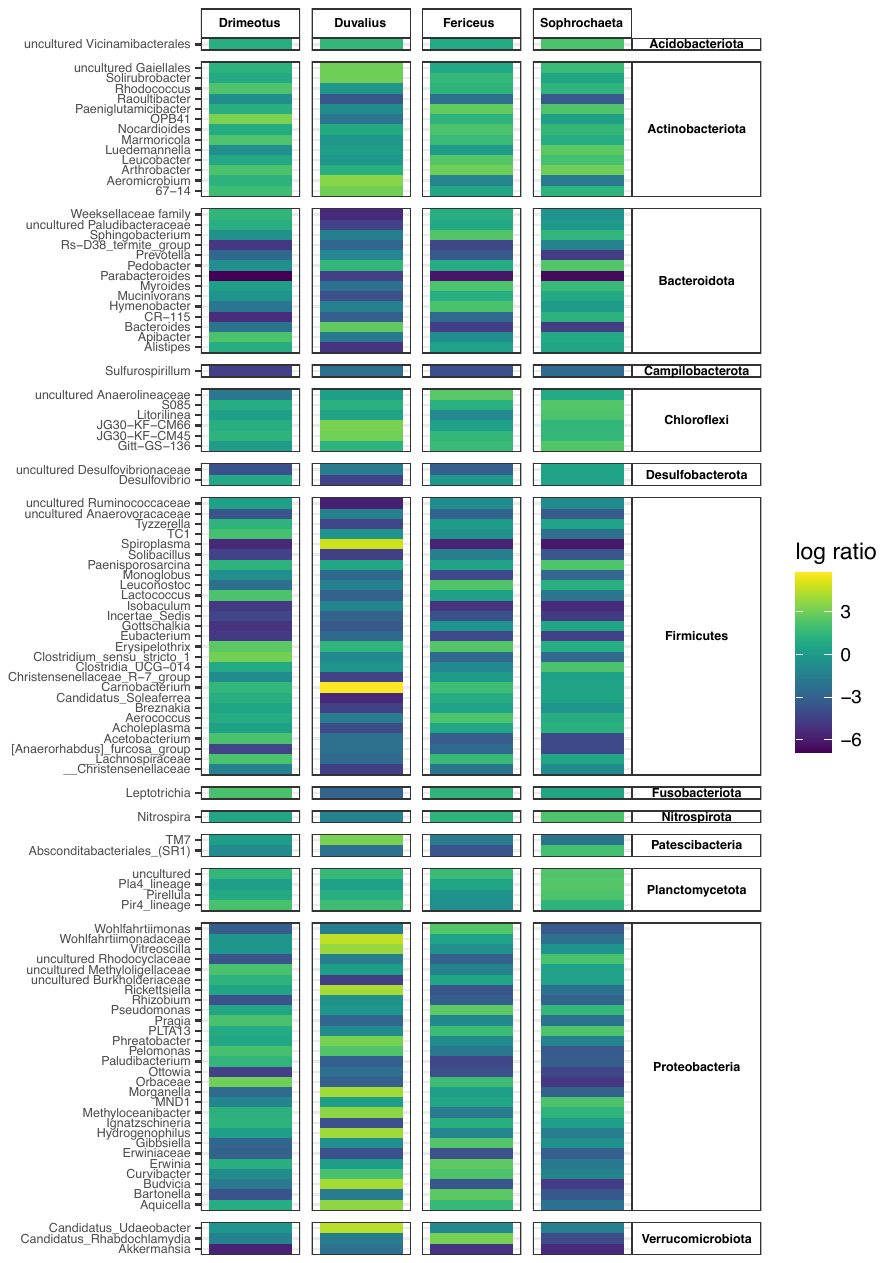

**Fig. S3:**  **Comparison between genera of the gut microbiomes in Coleoptera.** Heatmap of the bacteria distribution across the studied cave species using as reference taxa *Closania*. Only Coleoptera species with more individuals were considered in the analysis.

**Fig. S4:** **Distances between the gut microbiome of the different Coleoptera species.** Principal coordinate projection Bray Curtis dissimilarity also including phyla with lower diversity.

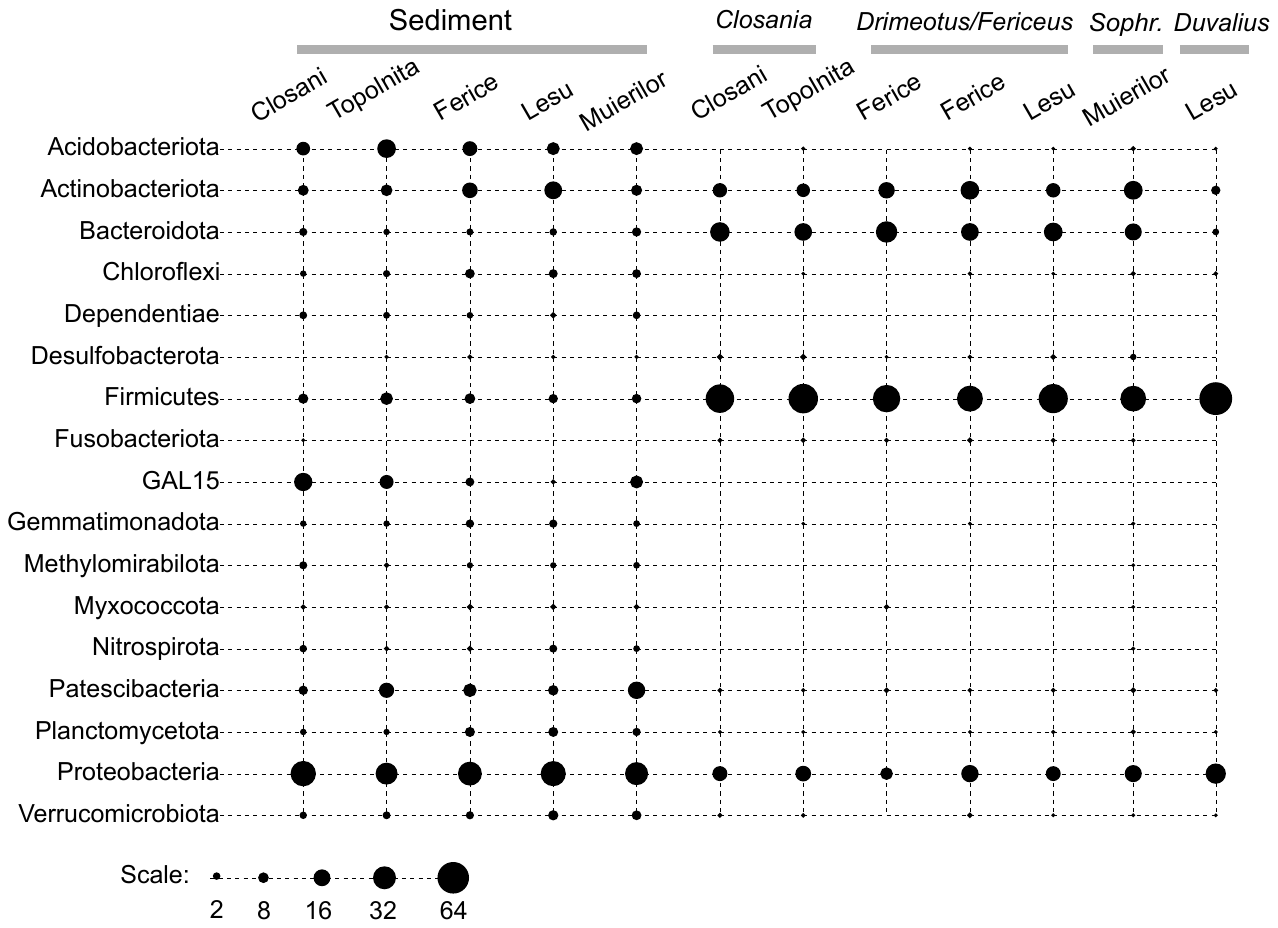

**Fig. S5:** **Distribution of phyla abundance across sediments and guts of the different studied caves and species.** Bubble plot representation.

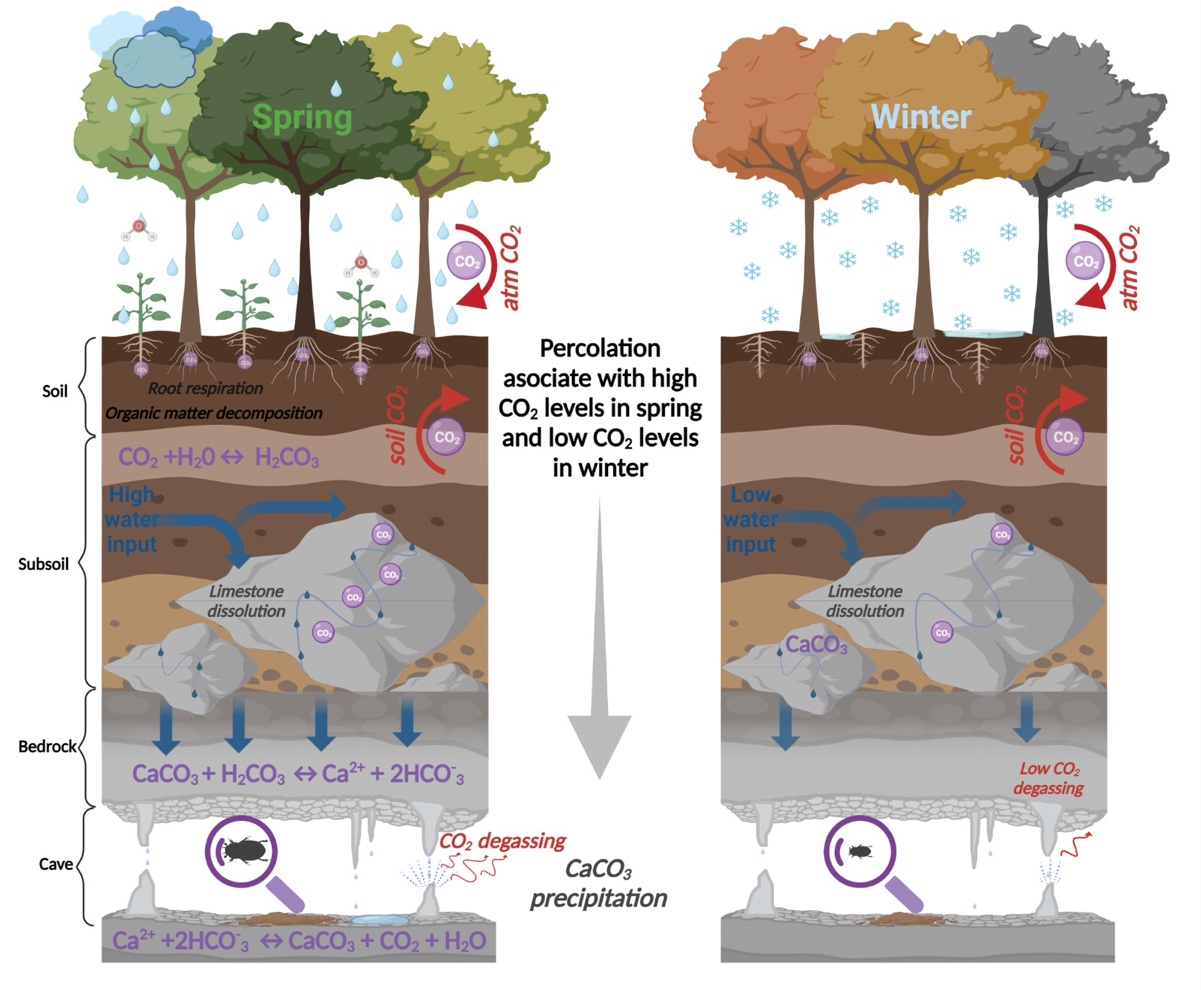

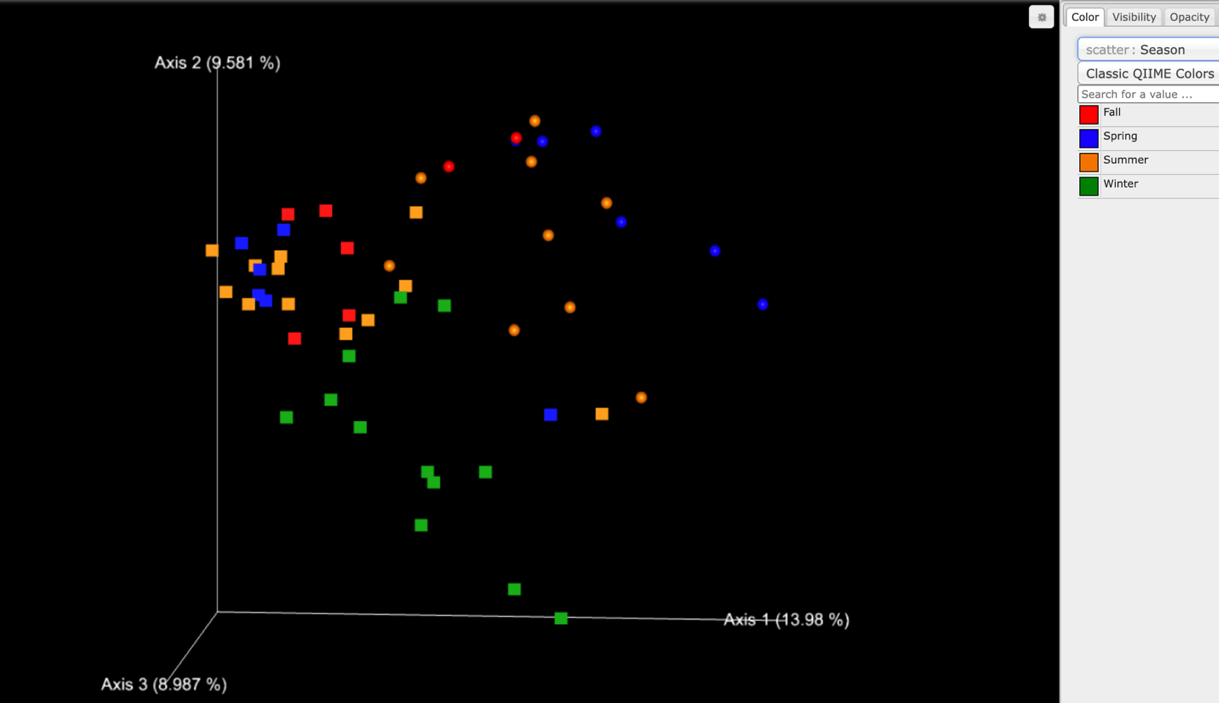

**Fig. S6:** **The seasonal variations in nutrients and the gut microbiome of *Drimeotus* from Lesu.** The seasonal variation of the surface nutrients input (above) and the seasonal distribution of the gut diversity (below) showing a larger variability for the gut microbiome of the individuals during spring (green squares).

**Table S1:** **The specimens of cave Coleoptera from which gut was isolated and analyzed for its microbiome.** *samples with analyzed sediment microbiome.

| **Cave (used name in the text)**  **Stations** | **Sample ID** | ***n*** | **Month**  **(2019)** | **Taxon** |
| --- | --- | --- | --- | --- |
| **Lesu**  Station L1  Station L1A  Station L2  Station L4 | L1IIID* | 6 | March | *Drimeotus viehmanni* (**D**) |
|  | L2IIID* | 6 |  |  |
|  | L1VD* | 3 | May |  |
|  | L2VD | 3 |  |  |
|  | L1VIIID* | 3 | August |  |
|  | L1AVIIID* | 3 |  |  |
|  | L2VIIID* | 3 |  |  |
|  | L4VIIID | 3 |  |  |
|  | L1XID* | 2 | November |  |
|  | L1AXID* | 3 |  |  |
|  | L4IIIDu* | 1 | March | *Duvalius* *paroecus* (**Du**) |
|  | L4VDu | 1 | May |  |
|  | L1VIIIDu* | 2 | August |  |
|  | L2VIIIDu* | 1 |  |  |
|  | L1XIDu* | 1 | November |  |
|  | L4XIDu* | 3 |  |  |
| **Ferice**  Station F1  Station F2  Station F3 | F1IIIF* | 3 | March | *Drimeotus (Fericeus) kraatzi* (**F**) |
|  | F2VF* | 3 | May |  |
|  | F1VIIIF* | 3 | August |  |
|  | F2VIIIF* | 3 |  |  |
|  | F3VIIIF* | 3 |  |  |
|  | F1XIF | 2 | November |  |
| **Closani**  Station C1  Station C3 | C1VCl* | 5 | May | *Closania winkleri* (**Cl**) |
|  | C3VCl | 3 |  |  |
|  | C1VIIICl | 3 | August |  |
|  | C3VIIICl | 3 |  |  |
|  | C1XICl | 2 | November |  |
| **Topolnita**  Station T1  Station T2  Station T3 | T1VCo* | 3 | May | *Closania orghidani* **(Co)** |
|  | T1VIIICo* | 3 | August |  |
|  | T2VIIICo | 2 |  |  |
|  | T3IXCo | 3 | November |  |
| **Muieri**  Station M2  Station M5  Station M11 | M2VS* | 3 | May | *Sophrochaeta chappuisi* **(S)** |
|  | M5VS* | 2 |  |  |
|  | M11VS | 2 |  |  |
|  | M2VIIIS* | 2 | August |  |
|  | M5VIIIS | 2 |  |  |
|  | M11VIIIS* | 2 |  |  |
|  | M11VDv* | 1 | May | *Duvalius voitestii* **(Dv)** |

**Table S2:** **The microclimatic conditions in the studied stations of the five caves** (see also Fig. S1 for the stations).

| Station | March | |  | May | |  | August | |  | November | |  | Substrate |
| --- | --- | --- | --- | --- | --- | --- | --- | --- | --- | --- | --- | --- | --- |
|  | **T (°C)** | **Rh (%)** | **CO_2_ (ppm)** | **T (°C)** | **Rh (%)** | **CO_2_ (ppm)** | **T (°C)** | **Rh (%)** | **CO_2_ (ppm)** | **T (°C)** | **Rh (%)** | **CO_2_ (ppm)** |  |
| L1 | 7.8 | 94 | 500 | 8.4 | 93 | 1400 | 11.1 | 71 | 1600 | 8.4 | 81 | 500 | Clay deposit, on the bank of the underground river |
| L1A | 8.2 | 97 |  | 8.7 | 96 |  | 11.3 | 85 |  | 9.0 | 90 |  | Gravel, washed, with a film of water |
| L2 | 8.6 | 91 |  | 8.8 | 90 |  | 9.3 | 86 |  | 8.5 | 82 |  | Cave wall with a water film, soaked clay at the base of the wall |
| L4 | 8.0 | 99 |  | 8.3 | 95 |  | 9.1 | 85 |  | 8.7 | 92 |  | Fine clay substrate at the base of the wall |
| F1 | 11.4 | 88 | 900 | 13.3 | 75 | 2000 | 12.7 | 77 | 3100 | 11.5 | 91 | 2100 | Clay on the cave floor |
| F2 | 12.2 | 88 |  | 13.0 | 79 |  | 11.8 | 88 |  | 11.4 | 93 |  | Clay on the cave floor |
| F3 | 11.9 | 90 |  | 12.7 | 86 |  | 12.7 | 86 |  | 11.4 | 96 |  | Clay on the cave floor |
| C1 | 12.3 | 92 | 800 | 11.2 | 89 | 1200 | 13.2 | 76 | 6600 | 11.3 | 96 | 2800 | Clay deposit near the cave wall |
| C3 | 12.4 | 91 |  | 11.6 | 91 |  | 12.6 | 83 |  | 11.2 | 95 |  | Stalagmite floor with water film |
| T1 | 11.7 | 95 | 1000 | 14.1 | 83 | 1300 | 11.3 | 94 | 2800 | 11.3 | 97 | 800 | Clay on the cave floor |
| T2 | 12.1 | 94 |  | 12.3 | 91 |  | 11.5 | 94 |  | 11.3 | 95 |  | Crust on the cave floor, with clay deposits |
| T3 | 12.3 | 94 |  | 12.6 | 88 |  | 11.7 | 94 |  | 11.4 | 95 |  | Sandy substrate on the cave floor |
| M2 | 12 | 89 | 600 | 10.7 | 99 | 500 | 12.5 | 75 | 600 | 10.3 | 93 | 600 | Gravels with yellow clay deposits |
| M5 | 10.5 | 93 |  | 9.2 | 98 |  | 10.6 | 85 |  | 9.7 | 93 |  | Hardened clay on the cave floor |
| M11 | 10.1 | 95 |  | 10 | 97 |  | 10.0 | 92 |  | 10.8 | 91 |  | Fine clay on the cave floor |

**Table S3:** **Mineralogical content of sediment samples in the five studied caves** (see also Fig. S1 for the stations).

|  | Quartz | Calcite | Chlorite | Muscovite | Albite | Dolomite | Apatite |
| --- | --- | --- | --- | --- | --- | --- | --- |
| L1 | 33 | 22 | 5 | 40 | 0 | 0 | 0 |
| L2 | 50 | 3 | 0 | 47 | 0 | 0 | 0 |
| L4 | 11.4 | 46.6 | 0 | 42 | 0 | 0 | 0 |
| F1* | 20-50% | 0 | 0 | 5-20% | <5% | 0 | 0 |
| F2 | 84.77 | 0 | 3.66 | 11.57 | 0 | 0 | 0 |
| F3 | 48.22 | 0 | 47.74 | 4.04 | 0 | 0 | 0 |
| C1 | 21 | 0 | 0 | 45 | 34 | 0 | 0 |
| C3 | 43.26 | 0 | 0 | 39.25 | 17.49 | 0 | 0 |
| T1 | 44.92 | 23.04 | 11.67 | 7 | 13.37 | 0 | 0 |
| T2 | 38.79 | 0 | 14.86 | 14.45 | 31.9 | 0 | 0 |
| T3 | 43.83 | 31.32 | 4.46 | 8.42 | 11.97 | 0 | 0 |
| M2 | 16 | 51 | 0 | 0 | 0 | 0 | 33 |
| M5 | 35 | 0 | 0 | 41 | 24 | 0 | 0 |
| M11 | 37.83 | 0 | 0 | 38.3 | 13.65 | 10.22 | 0 |

* a different method was used for this sample

**Table S4:** **Physico-chemical parameters of sediment samples** (see also Table S1). Measurement units are given in mg/kg except for pH (pH units), electrical conductivity (µS/cm), N (%), and C (%).

| **Element** | **Samples** | | | | | | | | | | | | | | | | | | | | | | |
| --- | --- | --- | --- | --- | --- | --- | --- | --- | --- | --- | --- | --- | --- | --- | --- | --- | --- | --- | --- | --- | --- | --- | --- |
|  | **Lmar** | **L1aug** | **L1nov** | **L1Anov** | **L2mar** | **L4mar** | **L4nov** | **F1mar** | **F1nov** | **C1may** | **C1aug** | **C1nov** | **T1may** | **T1aug** | **T2aug** | **T3nov** | **M2may** | **M2aug** | **M5may** | **M5aug** | **M5nov** | **M11may** | **M11aug** |
| **pH** | 7.6 | 8.5 | 7.8 | 8.6 | 8.0 | 8.3 | 8.0 | 8.4 | 8.8 | 8.1 | 7.9 | 8.6 | 8.1 | 8.5 | 8.5 | 9.0 | 9.0 | 8.0 | 8.2 | 8.3 | 8.5 | 8.0 | 8.4 |
| **EC** | 259 | 100 | 96.7 | 83.9 | 131 | 104 | 246 | 151 | 83.1 | 78.8 | 38.5 | 51.9 | 80.3 | 63.5 | 95.9 | 111 | 59.0 | 280 | 122 | 110 | 93.4 | 155 | 156 |
| **N** | <0.01 | <0.01 | 0.30 | 0.56 | 0.25 | <0.01 | 0.21 | <0.01 | <0.01 | <0.01 | 0.07 | <0.01 | <0.01 | <0.01 | <0.01 | <0.01 | <0.01 | 0.23 | 0.48 | 0.52 | 0.35 | 0.46 | 0.41 |
| **C** | 2.12 | 2.20 | 3.54 | 5.04 | 1.14 | 0.28 | 2.28 | 0.70 | 0.41 | <0.01 | <0.01 | <0.01 | <0.01 | 0.12 | 0.71 | 1.66 | 9.80 | 8.05 | 2.82 | 3.57 | 1.96 | 2.15 | 1.65 |
| **Na** | 130 | 453 | 75.3 | 123 | 254 | 67.5 | 123 | 186 | 115 | 477 | 540 | 124 | 347 | 583 | 543 | 171 | 440 | 2700 | 8117 | 620 | 119 | 331 | 837 |
| **Mg** | 3147 | 26567 | 24690 | 5134 | 1756 | 1166 | 4305 | 2957 | 15093 | 8290 | 7810 | 9707 | 7590 | 9123 | 7227 | 6820 | 4393 | 3283 | 2723 | 1147 | 1953 | 3617 | 2000 |
| **K** | 1466 | 863 | 1861 | 2193 | 4253 | 1452 | 1908 | 4533 | 2663 | 2973 | 3420 | 2333 | 2143 | 2963 | 2317 | 1807 | 1283 | 1417 | 3073 | 2117 | 2668 | 3743 | 2933 |
| **Ca** | 30527 | 61933 | 49333 | 110933 | 32333 | 22323 | 69400 | 32790 | 31067 | 17443 | 13463 | 16687 | 11697 | 12403 | 30257 | 113667 | 487333 | 341333 | 88000 | 100400 | 67333 | 126100 | 199900 |
| **Al** | 12283 | 10373 | 12057 | 19853 | 22800 | 10227 | 17993 | 20907 | 21433 | 39700 | 40600 | 26100 | 28147 | 33433 | 25153 | 14867 | 8013 | 6953 | 20463 | 12793 | 19353 | 25423 | 21283 |
| **Fe** | 7357 | 16917 | 17467 | 26813 | 8713 | 8463 | 28653 | 8450 | 30460 | 40167 | 43433 | 44340 | 30417 | 47967 | 30387 | 23100 | 6723 | 6037 | 14327 | 22720 | 30047 | 23123 | 19323 |
| **P** | 829 | 1292 | 658 | 4247 | 11723 | 1146 | 5240 | 3383 | 3913 | 2690 | 1326 | 1178 | 2847 | 1161 | 246 | 2022 | 4657 | 20893 | 17977 | 6460 | 14780 | 52033 | 40067 |
| **S** | 266 | 131 | 118 | 734 | 494 | 158 | 332 | 130 | 59.1 | 73.8 | 42.6 | 35.9 | 124 | 70.2 | 58.5 | 207 | 255 | 186 | 468 | 349 | 234 | 472 | 298 |
| **V** | 28.2 | 18.4 | 39.0 | 19.4 | 51.9 | 30.7 | 17.8 | 52.2 | 24.4 | 26.2 | 46.5 | 36.0 | 34.2 | 57.9 | 36.1 | 23.8 | 4.7 | 7.6 | 12.9 | 18.3 | 16.8 | 10.8 | 16.7 |
| **Cr** | 14.7 | 11.7 | 22.0 | 14.0 | 26.5 | 15.3 | 13.0 | 33.9 | 20.0 | 16.9 | 28.2 | 26.0 | 25.4 | 37.9 | 29.8 | 22.1 | 3.1 | 5.8 | 22.1 | 15.9 | 21.2 | 19.6 | 14.5 |
| **Mn** | 492 | 313 | 556 | 350 | 982 | 809 | 436 | 577 | 401 | 317 | 580 | 512 | 199 | 354 | 388 | 177 | 95.0 | 252 | 214 | 101 | 107 | 262 | 225 |
| **Co** | 4.9 | 4.0 | 6.9 | 4.0 | 7.1 | 12.3 | 4.7 | 7.7 | 5.7 | 4.2 | 8.6 | 12.4 | 3.9 | 10.1 | 7.8 | 5.1 | 1.0 | 1.7 | 1.9 | 1.7 | 2.4 | 2.8 | 2.4 |
| **Ni** | 10.6 | 11.4 | 17.9 | 14.3 | 22.5 | 26.9 | 15.9 | 20.8 | 17.2 | 8.3 | 19.0 | 21.5 | 7.8 | 16.1 | 16.8 | 9.1 | 11.3 | 18.9 | 7.5 | 9.8 | 5.0 | 11.2 | 17.0 |
| **Cu** | 15.6 | 4.2 | 7.5 | 49.9 | 131 | 28.7 | 55.8 | 39.7 | 49.7 | 7.91 | 17.6 | 11 | 5.06 | 12.9 | 11 | 24.9 | 2.9 | 38.6 | 197 | 287 | 581 | 162 | 131 |
| **Zn** | 78.4 | 15.8 | 45.7 | 139 | 399 | 113 | 170 | 309 | 207 | 37.4 | 41.0 | 36.0 | 13.9 | 30.1 | 23.9 | 68.0 | 13.0 | 175 | 130 | 70.9 | 286 | 143 | 200 |
| **As** | 8.7 | 8.0 | 2.3 | 16.5 | 26.7 | 45.4 | 20.2 | 5.8 | 4.6 | 6.5 | 4.1 | 2.1 | 0.98 | 3.1 | 2.6 | 1.3 | 1.3 | 1.8 | 2.1 | 3.6 | 2.6 | 2.5 | 4.4 |
| **Sr** | 18.3 | 13.3 | 18.3 | 22.0 | 35.2 | 15.4 | 19.5 | 38.6 | 19.7 | 26.2 | 56.7 | 18.3 | 13.8 | 30.0 | 20.0 | 17.7 | 41.1 | 117 | 35.5 | 56.7 | 67.9 | 37.8 | 117 |
| **Ba** | 46.0 | 15.8 | 41.7 | 52.5 | 119 | 59.1 | 57.2 | 103 | 69.3 | 66.4 | 78.8 | 117 | 41.9 | 64.4 | 46.8 | 39.6 | 9.3 | 25.9 | 104 | 70.4 | 172 | 70.4 | 44.5 |
| **La** | 9.3 | 8.4 | 24.1 | 11.8 | 17.3 | 22.5 | 10.0 | 13.0 | 9.3 | 13.2 | 13.8 | 15.0 | 9.53 | 11.8 | 8.07 | 3.8 | 1.26 | 1.31 | 10.9 | 6.6 | 8.8 | 9.59 | 5.3 |
| **Ce** | 16.7 | 16.0 | 47.4 | 20.5 | 29.3 | 29.8 | 18.6 | 27.0 | 22.0 | 33.7 | 36.1 | 37.4 | 18.7 | 26.7 | 19.7 | 8.9 | 2.51 | 2.53 | 17.3 | 13.8 | 19.8 | 15.0 | 11.3 |
| **Pb** | 7.4 | 9.9 | 8.2 | 12.0 | 11.7 | 19.1 | 12.0 | 30.0 | 20.0 | 8.2 | 13.4 | 12.0 | 5.9 | 8.7 | 8.7 | 4.0 | 38.4 | 21.1 | 5.0 | 11.7 | 12.0 | 3.8 | 7.2 |
